# Coral: Clear and customizable visualization of human kinome data

**DOI:** 10.1101/330274

**Authors:** Kathleen S. Metz, Erika M. Deoudes, Matthew E. Berginski, Ivan Jimenez-Ruiz, Bulent Arman Aksoy, Jeff Hammerbacher, Shawn M. Gomez, Douglas H. Phanstiel

## Abstract

Protein kinases represent one ofthe largest gene families in eukaryotes and play roles in a wide range of cell signaling processes and human diseases, making them attractive targets for drug development. The human kinome is extensively featured inhigh-throughput studiesgenerating genomic, proteomic, and kinase profiling data. Current tools for visualizing kinase data in the context of the human kinome superfamily are limited to encoding data through the addition of nodes to a low-resolution image of the kinome tree. We present Coral, a user-friendly interactive webapplication for visualizing both quantitative and qualitative data. Unlike previous tools, Coral can encode data in three features (node color, node size, and branchcolor), allowsthreemodesofkinomevisualization (the traditional kinome tree as well as radial and dynamic-force networks) and generates high-resolution scalable vector graphic files suitable for publication without the need for refinement in graphic editing software. Due to its user-friendly, interactive, and highly customizable design, Coral is broadly applicable to high-throughput studies of the human kinome. The source code and web application are available at github.com/dphansti/Coral. html and phanstiel-lab.med.unc.edu/Coral respectively.

## INTRODUCTION

The human kinome consists of over 500 different protein kinases, which regulate a broad range of cellular processes via substrate phosphorylation. These proteins have been implicated in a variety of diseases including pathological hypertrophy (Heineke and Molkentin, 2006; Vlahos et al., 2003), rheumatoid arthritis (Gaestel et al., 2009; Patterson et al., 2014), and cancer (Fleuren et al., 2016; Ventura and Nebreda, 2006). As a result, protein kinases are common candidates for drug targets and are increasingly the focus of large-scale studies (Cohen, 2002; Cohen and Alessi, 2013; Duong-Ly and Peterson, 2013; Hu et al., 2017; Li et al., 2016; Wu et al., 2016). High-throughput approaches to study kinase abundance and activity–including proteomics, genomics, and kinase profiling screens–have created a need to visualize and interpret results in the greater context of the human kinome.

Multiple paradigms have been established for representing kinome attributes. In 2002, Manning et al. generated a kinome map based on sequence similarity between publicly available and predicted protein kinase domains and categorized the 518 known human kinases into nine superfamilies (Manning et al., 2002). This dendrogram was then modified in reference to other trees and manually refined into an aesthetically pleasing image in which branches are colored by kinase group and kinase names are HTML links to websites with information specific to each kinase (www.cellsignal.com). Several groups have manually modified this image, encoding qualitative or quantitative kinase information in either branch color (Fleuren et al., 2016; Phanstiel et al., 2011) or in the color and size of circular nodes placed at the end of each branch (Hu et al., 2017; Li et al., 2016; Wu et al., 2016). These figures are powerful tools for data interpretation and communication but are labor intensive to create and often infeasible for large or more complex experimental data sets. To address this issue, several programs have been developed to automate kinome visualization including Kinome Render (Chartier et al., 2013), the NCGC Kinome Viewer (tripod.nih. gov), and KinMap (Eid et al., 2017), which have been widely adopted by the kinase research community. Because these existing methods encode information solely by adding nodes to the existing image created by Cell Signaling Technology, they suffer from three limitations: (1) they offer no ability to recolor kinase branches either for aesthetic purposes or to encode information, (2) they only feature a single paradigm for kinome representation, the semi-quantitative tree, and (3) they produce low-resolution images that are not well suited for publication.

Here we describe Coral, an interactive web application for kinome annotation which allows encoding of qualitative and quantitative kinase attributes in branch color, node color, and node size. Coral offers multiple modes of kinome visualization including the traditional kinome tree, as well as interactive static and dynamic network layouts enabled by the D3 javascript library. Importantly, the images produced by Coral are highly customizable and can be downloaded as high-resolution, publication quality vector images. Coral was designed to be user friendly and includes example data, extensive documentation, and example outputs for all features.

## RESULTS

Coral is a highly customizable kinome visualization tool that provides unprecedented flexibility in terms of attribute encoding and visualization modalities, allowing for the creation of high-resolution figures through a user-friendly platform.

Coral offers multiple modes of data input and supports a variety of kinase identifiers. The user can supply both qualitative and quantitative data by selecting from pre-populated pulldown menus or by pasting information from common data sources such as spreadsheet editors. Coral inputs and outputs are compatible with multiple identifiers including UniProt, ENSEMBL, Entrez and HGNC, as well as an alphanumeric version of the names displayed in the original kinome tree, referred to as their CoralID.

Data can be encoded in branch color, node color, node size, and any combination thereof. Coral supports categorical, sequential, and divergent color schemes by providing several color-blind friendly palettes, as well as the option to create your own. This allows for straightforward incorporation of multiple types of data within a single figure, using intuitive color schemes and automatically generated legends for ease of interpretation.

Coral features three modalities of visualization. First, through the use of a newly rendered scalable vector graphics (SVG) file, Coral outputs high-resolution images with data encoded in a kinome tree based on the one created by Cell Signaling Technology (**Figure 1A**). Second, data can be depicted in a circular radial kinase network with nodes for kinases, groups, families, and subfamilies (**Figure 1B**). Finally, kinase attributes can be displayed in a dynamic force-directed network that uses the same underlying network structure as the circle plot (**Figure 1C**). All plots can be downloaded as high-resolution vector graphic files suitable for publication without the need for further refinement in graphic editing software. In addition, Coral provides advanced options for customizing features including titles, legends, fonts, transparencies, and node highlights. Users can also browse a searchable and sortable table of kinases and their associated attributes.

**Figure 1:**
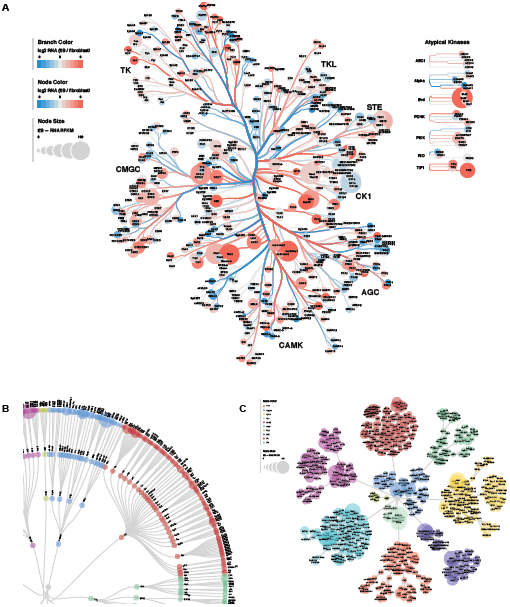
Three modes of kinome visualization. **(A)** A ‘Tree’ plot generated by Coral in which log2 RNA fold change in encoded in branch color and node color, and RNA RPKM values are encoded in node size. **(B)** A zoomed in image of a ‘Circle’ plot generated by Coral that depicts kinase group in node color and RNA RPKM values in node size. **(C)** A ‘Force’ plot generated by Coral depicting the same data as in panel B.

Coral was designed with user experience in mind. Each plot responds to user input, updates in real time, and features drag and zoom capabilities. Individual nodes are also interactive and either include links to UNIPROT pages or expand when hovered above making it easier to read text and visualize colors for individual nodes. Features are extensively documented in the Coral information pages which include both example data and example outputs for each feature to help familiarize users with its many options.

## DISCUSSION

Due to its flexibility and wealth of customizable features, Coral can potentially be applied to many areas of study within human kinome research. Quantitative data can be presented through branch color, node color, and node sizes and can feature sequential information (e.g. transcript or protein levels, kinase activity, or -log p-values) as well as divergent values (e.g. log2 fold-change). Qualitative data such as built-in kinase family assignments can easily differentiate between different classes of kinases and can also be applied to highlight kinases of interest such as targets of kinase inhibitors.

The traditional and network kinome structures can be used to highlight different features of data depending on the application. For example, the Tree layout may be useful for sharing data among audiences familiar with the historically used phylogenetic tree, while the network layouts are useful when there is a need to explicitly include family and subfamily delineations and highlight the hierarchy of the kinase classifications.

With the constantly growing wealth of human kinase studies, Coral fills a need for easily-generated, customizable, high-resolution images for the intuitive visualization of kinase data in the broader context of the human kinome.

## METHODS

Coral was developed using R and Javascript.

The R package shiny (Chang, et al., 2017) and its extensions shinydashboard (Chang and Ribeiro, 2017), shinyBS (Bailey, 2015) and shinyWidgets (Perrier and Meyer, 2018) were used for the web framework.

The R packages readr (Wickham, et al., 2017) and rsvg (Ooms, 2017) were used for data manipulation and rendering the SVG elements.

RColorBrewer (Neuwirth, 2014) was utilized for color palettes.

The Circle and Force layouts were written using the D3.js library (Bostock, et al., 2011).

The kinome tree was obtained courtesy of Cell Signaling Technology Inc. (www.cellsignal.com), and manually redrawn using vector graphics in Adobe Illustrator.

## AUTHOR CONTRIBUTIONS

Conceptualization: D.H.P.

Software: D.H.P., E.M.D., M.E.B, B.A.A., I.J.R., and K.S.M.

Writing: K.S.M. and D.H.P.

Visualization: E.M.D.

Supervision: D.H.P.

Resources: D.H.P., S.M.G, and J.H.

## ACKNOWLEDGEMENTS

We thank Danielle Swaney, Emily Cousins, and Craig Wenger for helpful conversations and software testing.

## REFERENCES

Bailey, E. 2015. shinyBS: Twitter Bootstrap Components for Shiny. Release R package version 0.61. https://CRAN.R-project.org/pack-age=shinyBS

Bostock, M., Ogievetsky, V. and Heer, J. D3: Data-Driven Documents. IEEE Transactions on Visualization and Computer Graphics 2011; 17(12):2301–2309.

Chang, W., et al. 2017. shiny: Web Application Framework for R. Release R package version 1.0.5. https://CRAN.R-project.org/pack-age=shiny

Chang, W. and Ribeiro, B.B. 2017. shinydashboard: Create Dashboards with ‘Shiny’. Release R package version 0.6.1. https://CRAN.R-project.org/package=shinydashboard

Chartier, M., et al. Kinome Render: a stand-alone and web-accessible tool to annotate the human protein kinome tree. PeerJ 2013;1:e126.

Cohen, P. Protein kinases--the major drug targets of the twenty-first century? Nat Rev Drug Discov 2002;1(4):309–315.

Cohen, P. and Alessi, D.R. Kinase drug discovery--what’s next in the field? ACS Chem Biol 2013;8(1):96–104.

Duong-Ly, K.C. and Peterson, J.R. The human kinome and kinase inhibition. Curr Protoc Pharmacol 2013;Chapter 2:Unit2 9.

Eid, S., et al. KinMap: a web-based tool for interactive navigation through human kinome data. BMC Bioinformatics 2017;18(1):16.

Fleuren, E.D., et al. The kinome ‘at large’ in cancer. Nat Rev Cancer 2016;16(2):83–98.

Gaestel, M., Kotlyarov, A. and Kracht, M. Targeting innate immunity protein kinase signalling in inflammation. Nat Rev Drug Discov 2009;8(6):480–499.

Heineke, J. and Molkentin, J.D. Regulation of cardiac hypertrophy by intracellular signalling pathways. Nat Rev Mol Cell Biol 2006;7(8):589– 600.

Hu, Y., Kunimoto, R. and Bajorath, J. Mapping of inhibitors and activity data to the human kinome and exploring promiscuity from a ligand and target perspective. Chemical Biology & Drug Design 2017;89(6):834–845.

Li, Y.H., et al. The Human Kinome Targeted by FDA Approved Multi-Target Drugs and Combination Products: A Comparative Study from the Drug-Target Interaction Network Perspective. PLoS One 2016;11(11):e0165737.

Manning, G., et al. The Protein Kinase Complement of the Human Genome. Science 2002;298:1912–1934.

Neuwirth, E. 2014. RColorBrewer: ColorBrewer Palettes. Release R package version 1.1-2. https://CRAN.R-project.org/package=RCol-orBrewer

Ooms, J. 2017. rsvg: Render SVG Images into PDF, PNG, PostScript, or Bitmap Arrays. Release R package version 1.1. https://CRAN.R-proj-ect.org/package=rsvg

Patterson, H., et al. Protein kinase inhibitors in the treatment of inflammatory and autoimmune diseases. Clin Exp Immunol 2014;176(1):1–10.

Perrier, V. and Meyer, F. 2018. shinyWidgets: Custom Inputs Widgets for Shiny. Release R package version 0.4.1. https://CRAN.R-project.org/package=shinyWidgets

Phanstiel, D.H., et al. Proteomic and phosphoproteomic comparison of human ES and iPS cells. Nat Methods 2011;8(10):821–827.

Ventura, J.J. and Nebreda, A.R. Protein kinases and phosphatases as therapeutic targets in cancer. Clin Transl Oncol 2006;8(3):153–160.

Vlahos, C.J., McDowell, S.A. and Clerk, A. Kinases as therapeutic targets for heart failure. Nat Rev Drug Discov 2003;2(2):99–113.

Wickham, H., Hester, J. and Francois, R. 2017. readr: Read Rectangular Text Data. Release R package version 1.1.1. https://CRAN.R-proj-ect.org/package=readr

Wu, P., Nielsen, T.E. and Clausen, M.H. Small-molecule kinase inhibitors: an analysis of FDA-approved drugs. Drug Discov Today 2016;21(1):5–10.

